# Response of land use change and ecosystem services based on different functional scenarios: Case study of Nanjing

**DOI:** 10.1101/2020.09.07.285726

**Authors:** Ziyi Wang, Yu wang, Jingxiang Zhang, Dongqi Sun, Zihang Zhou

## Abstract

Land Use/Land Cover Change (LUCC) is one of the important reasons for the change of ecosystem services (ESs). Due to the uncertainty of future development policies and the complexity of LUCC, assessing the impact of future urban sprawl on ecosystem services remains challenging. We simulated the effect of urban land-use change on ESs on the basis of different functional scenarios, which is of important value to urban land-use planning and ESs protection. In our study, we designed three scenarios: Production function priority scenario (PFP scenario)、 Living function priority scenario (LFP scenario)、 Ecological function priority scenario (EFP Scenario). And we used the GeoSOS-FLUS software to realize visualization. Based on invest model, we evaluated five types of ESs: carbon storage, warter yield, habitat quality, water purification and soil conservation. Research showed that from 2000 to 2015, carbon storage, habitat quality and water production in Nanjing decreased significantly, soil conservation increased slightly, and the performance of the two indicators for water purification was not consistent. From different scenarios, carbon storage and habitat quality were the highest in EFP scenario, water yield was the highest in PFP scenario and soil conservation was the highest in LFP scenario. We analyzed the trade-offs among various ESs, found that the change of land-use types in cities does not fundamentally change the trade-offs among various ESs. We believed that the determination of the main function of LUCC was the first condition to judge the applicability of scenario, and the scenario simulation which integrated the main function of the city could provide more references for the related research.

## Introduction

Ecosystem Services (ESs) refer to the natural environmental conditions and functions formed and maintained by the ecosystem and human survival processes, including the provision services, regulatory services, culture services and support services [1]. ESs have redefined the relationship between human and environment in cities, and the contribution of nature protection and ecological environment to solving urban problems [2]. ESs has become the frontier and hotspot of geography, ecology and related disciplines [3,4]. The assessment of ESs reflects the complex relationship between human society and ecosystem, and explain primely how human behavior affects the function of ESs.

Land Use/Land Cover Change (LUCC) and ESs affect each other and restrict each other. LUCC changes the land cover status and affects the regional ecological process, which makes the structure and function of ecosystem change, and then causes ESs function’s change. The global assessment report of the Intergovernmental Scientific-Policy Platform on Biodiversity and Ecosystem Services (IPBES) shows that three quarters of global terrestrial environment and 66 per cent of marine area have been affected or transformed by human beings [4], and the loss of ESs associated with land use change is very serious, and it also brings challenges to the ecosystem environment in the future [5]. From the perspective of urban planning, it will be the new direction of researchers in related fields to realize the optimal allocation of ESs based on the combination of various land use types [6-8].

At present, researchers at home and abroad have discussed how urban expansion affects ESs from different scales. For example, on a global scale, Costanza estimated the economic value of global ESs, and concluded that global LUCC between 1997 and 2011 had resulted in a loss of ESs of between $4.3 and $20.2 trillion yr [9]. Goldstein et al. argued that globally, ecosystems affected by land-use decisions contain at least 260 GT of irrecoverable carbon [10],which must be protected through a range of expanded policies; at the national level. Ouyang et al [11]. found that China’s ESs, including food production, carbon sequestration, soil conservation, flood control, sand control and water conservation, had increased from2000 to 2010, and that only a slight decline in the supply of habitat quality. Wei et al [12]. found that Ecosystem Service Value (ESV) changes in China from 2000 to 2008 were relatively mild compared with the rest of the world over the same period, which may be attributed to the famous conversion of cropland to forests in China. On the regional scale, Hu et al [13].studied the response of Pearl River Delta ESV to LUCC, and concluded that it is of great significance to construct a mathematical model to reveal the ESV of urban agglomeration. Msofe assessed changes in ESV as a result of LUCC in the Kilombero region of Tanzania from 1990 to 2016 [14]. The existing studies on the relationship between urban expansion and ESV mainly focus on the correlation between the two, and only few studies that can quantify the loss intensity of ESs caused by urban expansion.

The uncertainties in the future also give rise to researchers’ concerns that the impacts of urban sprawl on ES could long-lasting for decades [15].The analysis of the past and future ESs is of great importance to understand the changes of global and regional ESs, and it also helps to better combine public policies to serve the urban construction with diverse functions of ESs. Ye et al [16].assessed and predicted LUCC and ESs of Kunshan from 2006 to 2030, and quantified spatial-temporal change in ESs based on assessment. Wang et al[17].combined multi-objective planning and Dyna-CLUE model, predicted the situation of LUCC in Wuhan in 2030, and studied the potential impacts of LUCC on ESs from a spatial perspective. Based on the relationship between urban expansion and its driving factors, Chen et al. combined Cellular Automaton (CA) and Geographically weighted regression (GWR) to predict Chongqing urban expansion and ESV loss in 2030[18]. However, most of the researches focus on the prediction of land use structure, and few studies are based on the different functions of land use to predict the development trend of urbanization and reveal its trade-off effect on ESs. [19].

The study of ESs is often associated with management decision-making processes to improve the effectiveness of ecosystems for human well-being [20,21]. LUCC has the comprehensive function of ecology, production and life, and the important performance of land use transformation is that the limited land resources are redistributed in quantity and space among various main functions. Based on the three types of land use main function classification system, the regional land use transformation can be connected with the regional function transformation development, which could be a new perspective to study the land use transformation. We chose Nanjing, a developed city in southeast China, as a case to study how LUCC under different functional scenarios will affect the supply of ESs. In this paper, we designed three main land use scenarios, Production function priority scenario (PFP scenario)、 Living function priority scenario (LFP scenario) and Ecological function priority scenario (EFP Scenario),and combined the Markov model with GeoSOS-FLUS platform to predict and visualize future LUCC. We then chose the InVEST model developed by the Natural Capital Project Team from Stanford University, which has been used to assess the amount of ESs and to guide ecosystem management and decision-making[22-25]. Specifically, we focused on the following three main goals: (1) dynamic modeling of current and future ES based on different dominant land use functions -- carbon storage (CS), habitat quality (HQ), water yield (WY), water purification (WP), and soil conservation (SC);(2)quantify the impact of LUCC on these ESs;(3) provide appropriate policy suggestions to support reasonable land-use strategies, and to optimize ESs management in our research area.

## Materials and Methods

### Study area

Nanjing (31°14 “-32°37” N, 118°22 “-119°14” E) is located in the eastern of China, the southern of Jiangsu and the middle part of the lower reaches of the Yangtze River. It has 11 districts under its jurisdiction (Fig 1). Since the beginning of the 21st century, Nanjing has undergone a rapid process of urbanization. The permanent population reached 8.27 million at the end of 2016. The total rural population stood at 1.9912 million. The urbanization level had risen to 82%. And the regional GDP at the end of 2016 exceeded 1 trillion yuan. Nanjing is located in the tropical monsoon climate of North Asia, with zonal vegetation of deciduous broad-leaved and evergreen Temperate broadleaf and mixed forest, and rich in animal and plant resources, and has a wide variety of ecosystem types. The green ecosystem is distributed along the mountains and hills in Jiangning, Lishui and Pukou District. Besides the Yangtze River, the wetland ecosystem is mainly distributed in Gaochun and Lishui District. Nanjing is one of the four garden cities in China, as well as one of the Historical capitals of China, which received a special honorary award from the United Nations Human Settlements Programme (UN-Habitat). In the new round of Urban Master Plan of Nanjing, the intensive land use pattern has been put forward, the new path of ecological priority green development should be actively explored, and the new pattern of harmonious development between man and nature also should be promoted.

**Fig. 1.**
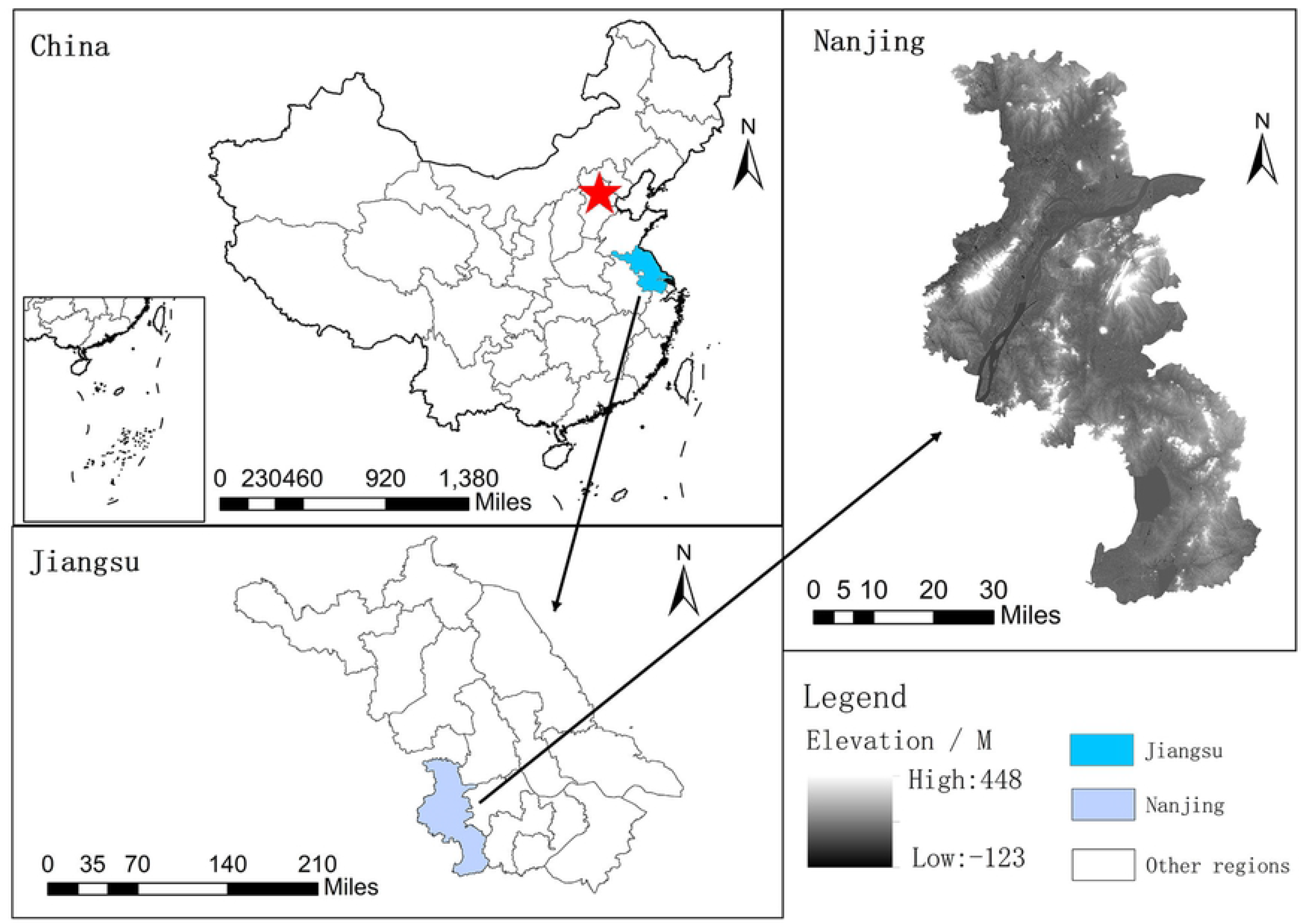
Study area.

### Future Land use modelling

#### Land use demand projection

Land use data for different policy scenarios in Nanjing are based on the GeoSOS-FLUS platform for future LUCC simulation, the model is a multi-scenario Simulation software based on the Future Land-Use Simulation developed by Liu Xiaoping and Li Xia of the National Sun Yatsen University. It is suitable for the simulation study of future LUCC scenarios. The model uses neural network (Ann) Algorithm to obtain the probability of suitability of all types of land, then coupling Markov Chain and cellular Automata (CA) model to improve the applicability of the model. In the CA model, an adaptive inertial competition mechanism is introduced to deal with the complexity and uncertainty of the mutual transformation of various land use types under the joint influence of nature and human activities.

##### (1) Land use demands in different functional scenarios

The paper is based on three functional scenarios: Production function priority scenario (PFP scenario)、 Living function priority scenario (LFP scenario)、 Ecological function priority scenario (EFP scenario). PFP scenario mainly considers the priority of protecting agricultural production space and limiting the transformation of agricultural land type to other land use types, while industrial production space and service industry production space are not the contents of this study. Life function is a place for people’s daily living and public activities, including urban living space and rural living space. Ecological function is the space to provide ecological products and services for the city and maintain the environmental quality. It is the natural base of production and living space. The EFP scenario requires that the ecological land types (woodland, grassland, water area) should be limited to other types. The simulation of the three scenarios needs to determine the Land-use demand as the input variable, and Land-use demands are based on the Markoff model to achieve the basic forecast and the combination of expert advice and relevant planning to determine.

##### (2) Visualization of LUCC in different functional scenarios

Firstly, we input the land use data of Nanjing City in 2000, and select the sampling mode of neural network training samples. According to our experience, we use uniform sampling mode to set the sampling parameter to 10 and the hidden layer number of neural network to 12. Secondly, we input 15 driving factors of LUCC, and standardize them [26-27].Then we set up three steps to achieve a more accurate multi-scenario land-use classification mapping:(1) Produce and input constraints on land-use change. The data are binary, with a value of 0 indicates that conversion of land types is not allowed in the area and 1 indicates that conversion is allowed. (2) Set up the cost matrix of land-use types in analog conversion. When one type of land-use does not allow conversion to another, we stipulate the relevant value of the matrix to 0; when conversion is allowed, we stipulate the relevant value of the matrix to 1. (Table S1) (3) Set the parameters of the neighborhood factors of land use types according to different scenarios. The parameter range is 0 ~ 1. The closer to 1, the stronger the expansion ability of land-use type is (Table S2).

#### Model implementation and precision validation

We tested and compared the performance of GeoSOS-FLUS model by LUCC from 2000 to 2015. In this study, the actual land-use cover map in 2015 was used as the control sample, and the forecast map in 2015 was used as the experimental sample. The closer the kappa coefficient is to 1, the better the simulation accuracy will be. When the Kappa Coefficient is greater than 0.8, which shows that the simulation accuracy of the model is satisfactory in statistical sense [28]. The result showed that the overall accuracy of the verification was 0.923, the Kappa Coefficient was 0.877, and the experimental simulation accuracy reached a high level. The result indicated that GeoSOS-FLUS model had a good applicability. Therefore, we chose the effective parameters and land-use classification map of 2015 to predict land-use classification maps of three scenarios in 2030.

#### Quantifying ESs

Invest version 3.8.0 was selected for the quantification of ESs. We focus on the following ESs: carbon storage, soil conservation, water yield, habitat quality, water purification. The data for each parameter was based on the published shared database.

#### Carbon Storage

The Invest Carbon model was used to quantify Carbon stocks in the study area. The model aggregates the amount of carbon stored in four different pools: aboveground biomass, underground biomass, soil carbon pool, dead organic matter [29](Table S3). The model requires a grid data set of current and future land-use maps which developed from scenario and biophysical data (including four carbon pools for each land use). The carbon pools of different land use types in Nanjing mainly come from previous studies [30-32].

We calculated the total carbon stored *CS_jxy_* for each given grid cell (x,y) with land use type ‘j’as:

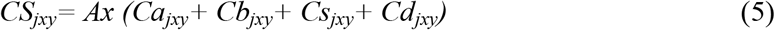

where *Ca_jxy_ Cb_jxy_, Cs_jxy_*, and *Cd_jxy_* are the carbon densities in the four main carbon pools of grid cell (x, y) in different land use types (j)

#### Water Yield

Water Yield (WY) refers to the ability of a piece of land to retain rainwater runoff. The calculation of water production service is based on the water balance method, and the data required for the calculation of water production service in this paper is calculated according to the relevant modules of invest model. The specific calculation formula is as follows:

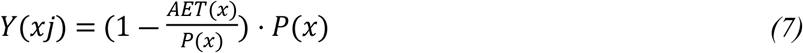

where Y(xj) is the annual water yield on grid x of different land use types, P(x) is the annual precipitation of grid x, and AET(xj)/P(x) is the vegetation evapotranspiration of different land use types [33-34].

#### Soil conservation

The soil conservation module uses the universal soil loss equation (USLE) to calculate potential soil loss and sediment transport in pixels for each patch, the module links the integrated use of land use and cover information, soil properties, digital elevation models of the study area, rainfall and meteorological and other data, and decision makers can influence soil erosion rates by changing land use and cover. The sediment retention model can be used to calculate the average annual soil loss per site, to determine the amount of sediment reaching the target site, to estimate the soil retention capacity of each site, and to estimate the cost of sediment removal(Table S5). The quality change of soil and water conservation can be judged by calculating the annual soil loss change of each pixel, the formula is as follows:

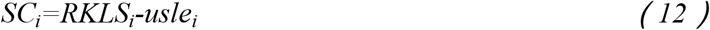

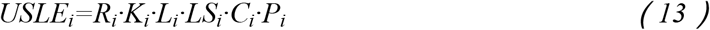

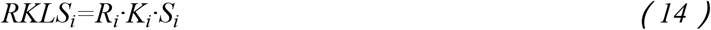

where *SC_i_* is the annual soil conservation amount of Nanjing; *USLE_i_* is the actual soil loss amount of Nanjing in pixel *i*; *R_i_* is the rainfall erosivity factor; *K_i_* is the soil erodibility factor; *R_i_* and *K_i_* represent the rainfall erosivity factor and the soil erodibility factor respectively; In addition, *LS_i_,S_i_,C_i_* and *P_i_* represent length gradient factor, slope factor, crop management and support practice factor respectively[26].

#### Water purification

The Nutrient Delivery Ratio Model (NDR) (Table S6) reflects the water purification degree of the watershed by estimating the amount of N and P nutrients retained by vegetation and soil in runoff. The greater the amount of N and P retained, the better the water purification service. The calculation is divided into two layers. First, the annual average runoff is calculated by the water production model, and the calculation process is the same as the water conservation module. Second, calculate the nutrient retention of each plaque:

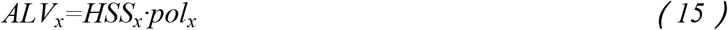

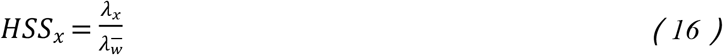

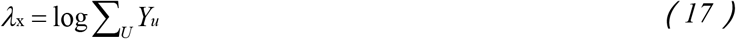

where *ALV_x_* represents the output nitrogen and phosphorus of the adjusted x grid cell,*pol_x_* and *HS S_x_* represents the outlet coefficient and hydrological sensitivity score of grid x respectively; *λ*x*λ* x is the runoff index of grid x, 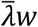 is the average runoff index of drainage basin, and *∑_U_∑_u_* is the sum of the water production of the grid cells along the flow path above the grid element x. [35,36].

#### Habitat quality

Habitat quality has been regarded as one of the significant ESs indicator [37]. Habitat quality module quantitatively assesses habitats spatially through interactions between habitats and their stressors(Table S7). Then the habitat distribution and degradation under different landscape patterns will be evaluated according to the response degree of different habitats to the stressors, so as to graphically represent the biodiversity index of the target lands. The habitat type, the stress intensity, the space distance of the habitat and the stress source, the sensitivity of the habitat to the stress source and others are the factors that affect the habitat quality. Based on previous studies,the impact *i_rxy_* of threat r from grid cell y on the habitat in grid cell x is represented by the following equations [38]:

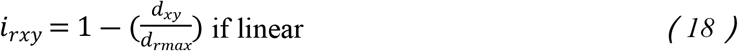

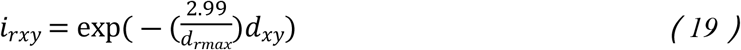

where *d_xy_* represents the straight line distance between grid x and y, and *d_rmax_* represents the maximum influence distance on stress factor r [39,40].

#### Assessment of the Trade-offs/Synergies among ESs

Due to the different responses of ESs to LUCC, it is the premise to realize the multi-objective protection and management of ESs to scientifically understand the trade-off and collaborative relationship between ESs. [41]. Therefore, it is necessary to represent the trade-off/collaborative relationship among various types of ESSS based on correlation coefficient. we use the “create random points” module in arcgis10.3 to select random sampling points for each scenario We use the “extract multiple values to points” module to extract the ES value of each sample point. In this study, In this study, 1000 samples were taken from each type of ESs. In the correlation analysis, we use Pearson parameters correlation test and draw based on Python 3.7. [42,43].

#### Data requirement and preparation

The 2000 and 2015 Nanjing rasterized land use maps (with a spatial resolution of 30 m) were obtained from the Department of Resources and Environmental Sciences of the Chinese Academy of Sciences (http://www.resdc.cn/). The data was based on the supervised classification of Landsat TM image by ENVI and was generated by manual visual interpretation. Six land categories are used in this paper: cropland, woodland, grassland, water area, build-up area and unused land. In future land-use model section, we used 15 driving factors from three categories (Fig.2): (1) physical condition, which includes digital elevation model, slope, undulation, organic matter, soil erosion, silt content, sand content, clay content, and precipitation; (2) spatial accessibility, which includes distances to residence, river, railways, roads and traffic ports; (3) socioeconomic factors, includes GDP and population. In the InVEST Model, in addition to the data mentioned above, we supplement the data by classification based on the needs of the evaluation of different ESs: (1) data obtained from the National Earth System Science Data Centre: Soil erosion data、 Plant Available Water Content、 Rainfall Erosion Index; (2) soil properties and depth of root limiting layers derived from the Harmonized World Soil Database (HWSD). (3) evapotranspiration coefficient (*K*c) of crops obtained from the Food and Agriculture Organization of the United Nations. (4) watershed scope data which obtained from CAS Resource and Environment Data Cloud Platform.

**Fig. 2.**
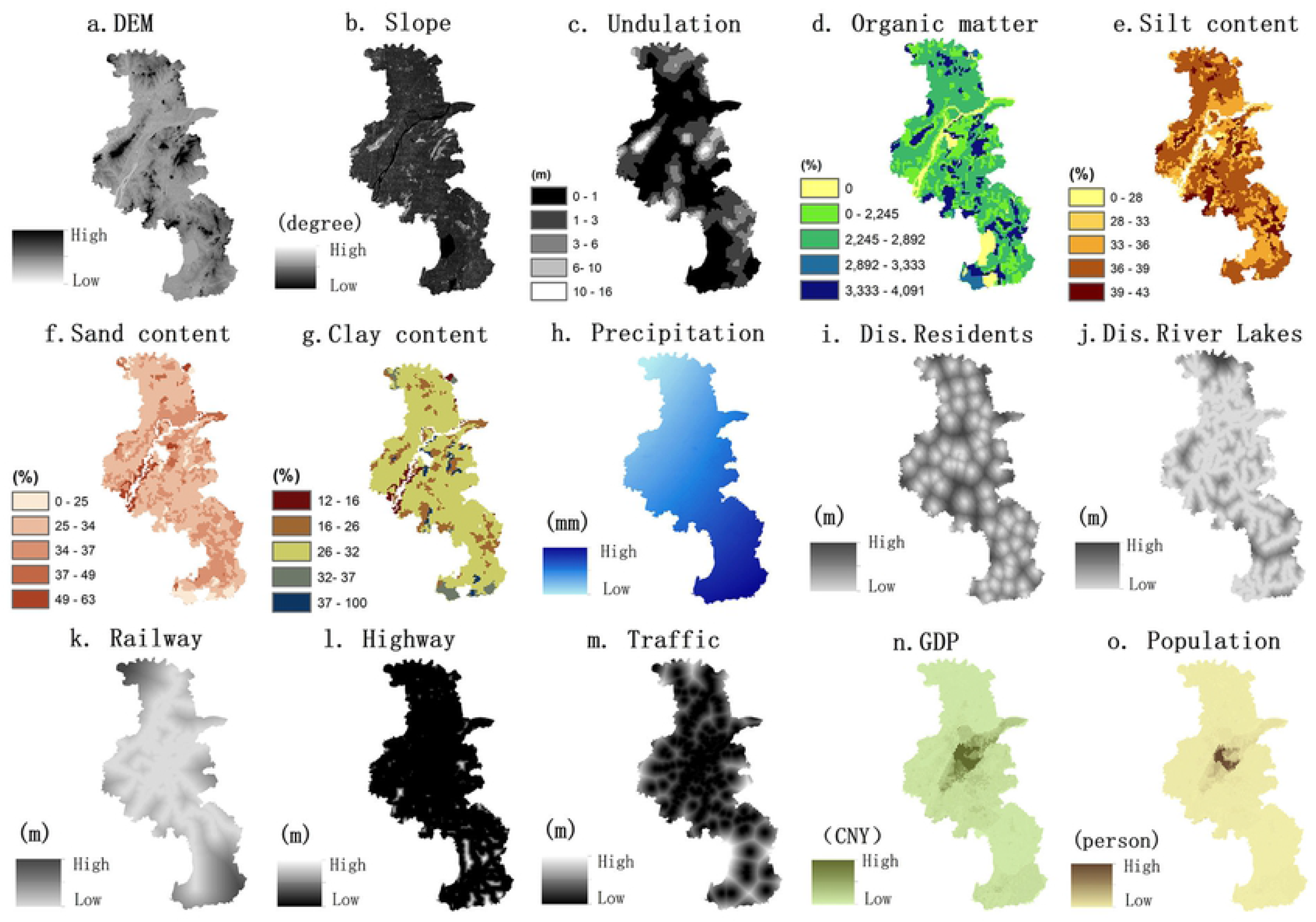
Driving factors of land use change in Nanjing.

## Results

### Actual and projected LUCC in Nanjing from 2000 to 2030

As shown in Nanjing land-use transfer Matrix in 2000-2015(Table 1, Fig. S1). The main land use types in Nanjing in 2015 were cropland, townland and water area, accounting for 54.33%, 23.96% and 10.3% respectively, the second land use types were woodland and grassland (10.23% and 0.83%), and with little unused land (0.35%). The trend of build-up land was in growth in the PFP、 LFP and EFP scenario(Fig.3), and the ratio of build-up land was expected to be 3.6%, 6.3% and 2.4%. while the proportion of cropland was expected to decrease by 3.1%, 5.9% and 4.7%. Meanwhile the types of land that were expected to decrease also include water area and unused land. The proportion of woodland in the PFP scenario was decreased, while that in the LFP and EFP scenarios was increased. Based on the horizontal comparison of different land use types in different scenarios in 2015, Build-up land raised by 26% in LFP scenarios, which was the largest increase among the three scenarios. The main reason was that the development orientation of living space needed to relax the restriction of urban expansion relatively. The area of cropland was in decline in all scenarios, but the decrease was minimal in the PFP scenario. The area of Woodland and Grassland both increased more than 20% in the EFP scenario. The change range of Water area and Unused land were small, but both are in the trend of area reduction.

**Table 1.**
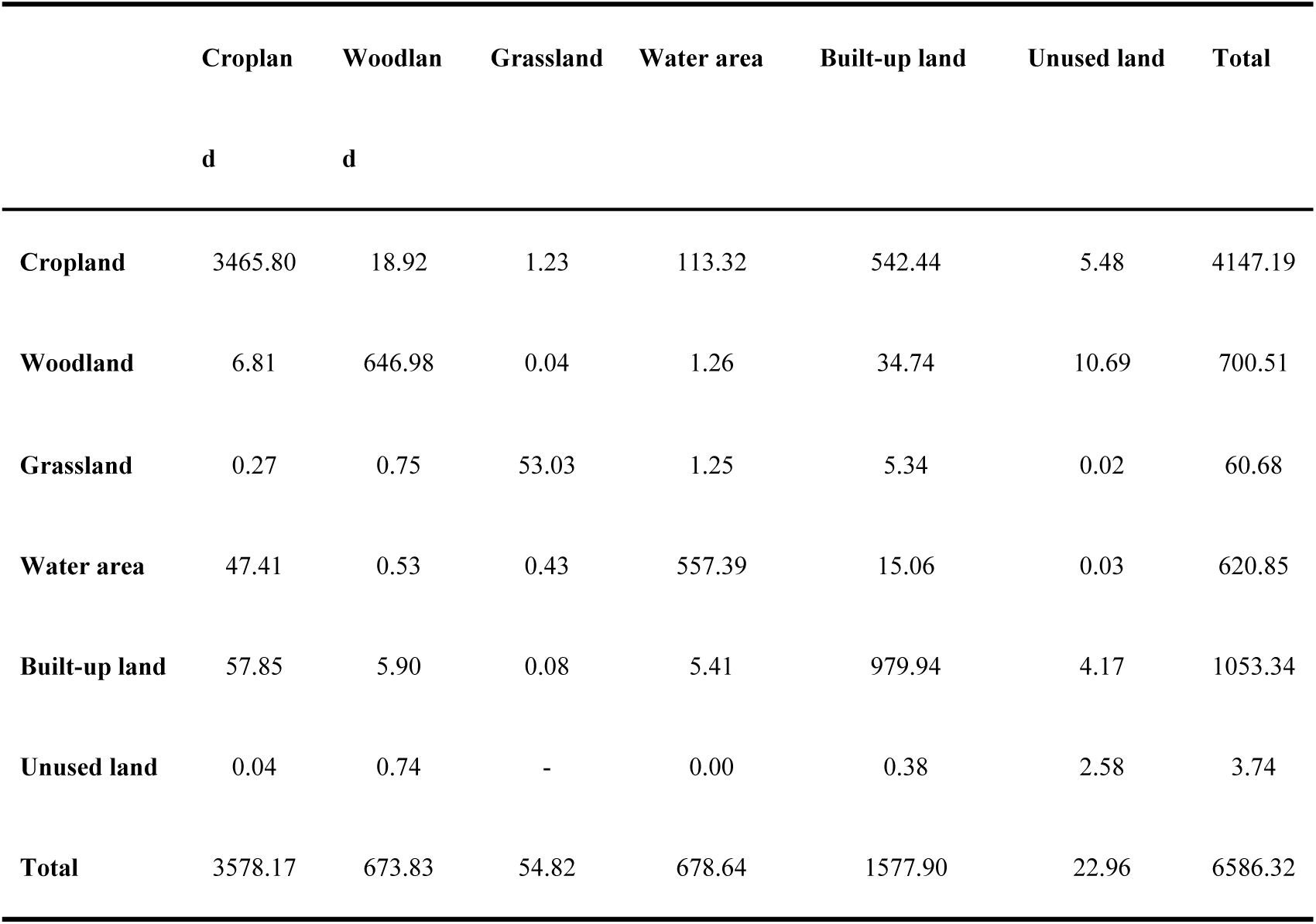
Land use transfer matrix of Nanjing from 2000 to 2015 (Km^2^)

**Table 2.** ESs change matrix driven by per-unit land use transitions of the main land use types from 2015 to 2030 under the PFP, LFP, and EFP scenarios.

|  | Conversions | Area (km<sup>2</sup>) | CS<br>(10<sup>4</sup> Mg) | WY<br>(10<sup>6</sup> m<sup>3</sup>) | SC<br>(10<sup>5</sup> t) | HQ<br>(unitless) | PL<br>(10<sup>3</sup>Kg) | NL<br>(10<sup>4</sup> Kg) |
| --- | --- | --- | --- | --- | --- | --- | --- | --- |
|  | 2—1 | 12.35 | -3.79 | 4.08 | -12.95 | -3955.08 | 0.63 | 0.19 |
| **2015-2030** | 2—5 | 6.30 | -2.61 | 4.32 | 0.22 | -3457.7 | 0.017 | 0.17 |
| **(PFP)** | 4—1 | 5.53 | 0.26 | 0.15 | 1.54 | 785.2 | 0.19 | 0.052 |
|  | 6—5 | 3.78 | 2.43 | 2.04 | 3.63 | 56.47 | -0.020 | -0.024 |
|  | 1—4 | 41.06 | -5.91 | -36751.8 | -59.47 | 16821.6 | -4.37 | -1.13 |
| **2015-2030** | 1—5 | 407.61 | 102.91 | 242.89 | 410.94 | -207203.7 | -51.77 | 11.28 |
| **(LFP)** | 4—5 | 5.20 | 14.58 | 1.54 | 0.14 | -74.48 | -0.008 | 0.091 |
|  | 6—2 | 11.11 | -0.19 | -7.07 | 19.6 | 11393.15 | -0.84 | -0.43 |
|  | 1—2 | 157.56 | 80.35 | -86.91 | 0.56 | 84277.89 | -17.45 | -4.47 |
| **2015-2030** | 1—4 | 71.19 | -12.81 | -97075.5 | 0.039 | -36509.87 | -9.63 | -2.54 |
| **(EFP)** | 1—5 | 148.31 | -36.76 | 88.91 | 0.16 | -73663.85 | -18.53 | 3.98 |
|  | 2—5 | 17.53 | -12.81 | 20.93 | 0.073 | -17058.62 | 0.10 | 0.83 |

**Fig. 3.**
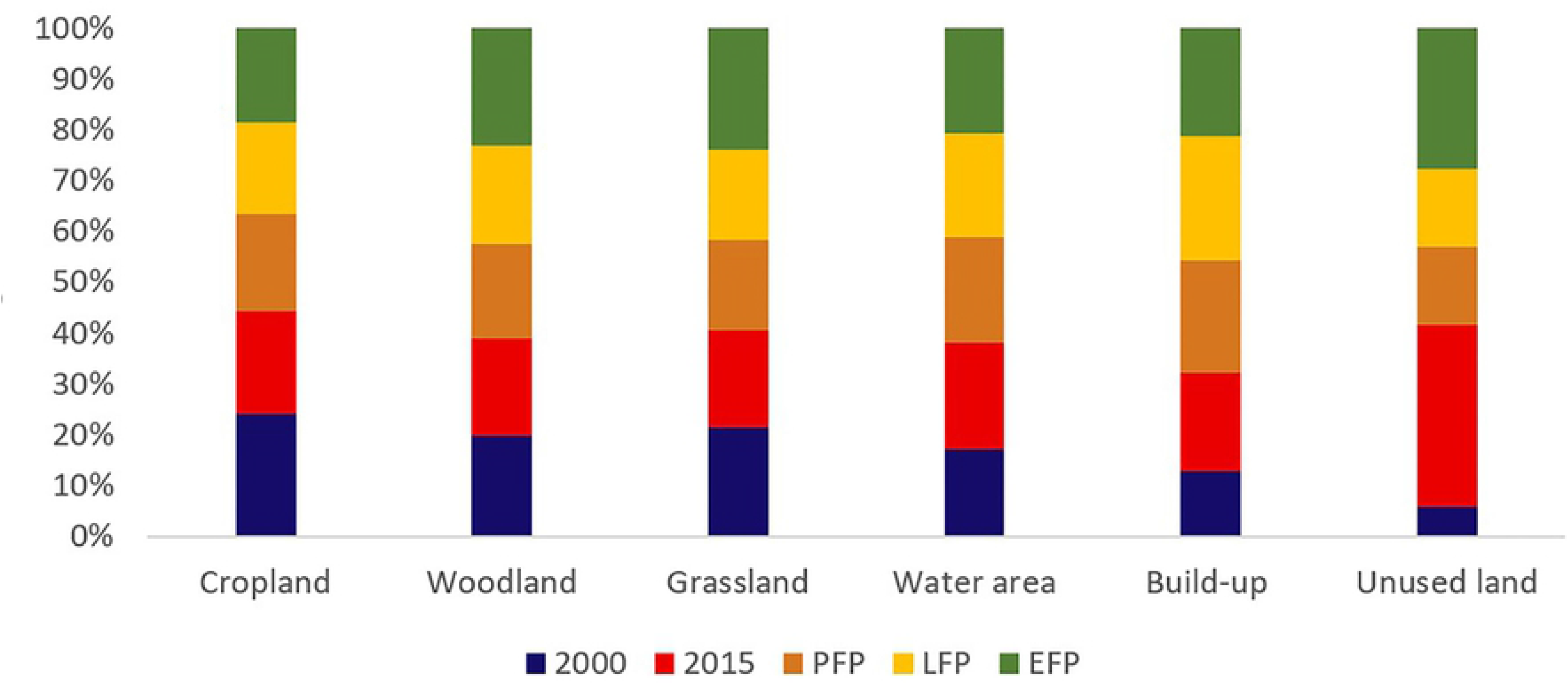
Actual and projected LUCC between 2000 to 2030 in Nanjing.

### ESs changes and tradeoff under different scenarios

As can be seen from figures 4, from 2000 to 2015, Nanjing’s carbon storage, habitat quality and water yield capacity declined by 17%, 21%, 20% respectively, and its soil and water conservation capacity still remained stable. From the perspective of different scenarios, ESs have different changes. Carbon storage in the PFP and LFP scenarios was projected to decrease by 3% and 5%, respectively from 2015 to 2030, while carbon storage in the EFP scenario was projected to increase by 8%; similarly, in the LFP scenario, habitat quality is expected to decline by 8%, and rise by 7%, 11% in the PFP and EFP scenarios. In the PFP scenario, the water yield capacity remained stable, while in the LFP and EFP scenarios it decreased by 2% and 16% respectively. In all three scenarios, the soil conservation capacity kept the trend of steady growth. Then we used phosphorus and nitrogen loads to characterize the input of pollutants. In PFP and LFP scenarios, from 2015 to 2030, the total nitrogen load in Nanjing is expected to show an upward trend, while the total phosphorus load to keep steady. In the EFP scenario, we expected that total nitrogen and total phosphorus loads to significantly reduced by 16% and 30% respectively.

**Fig. 4.**
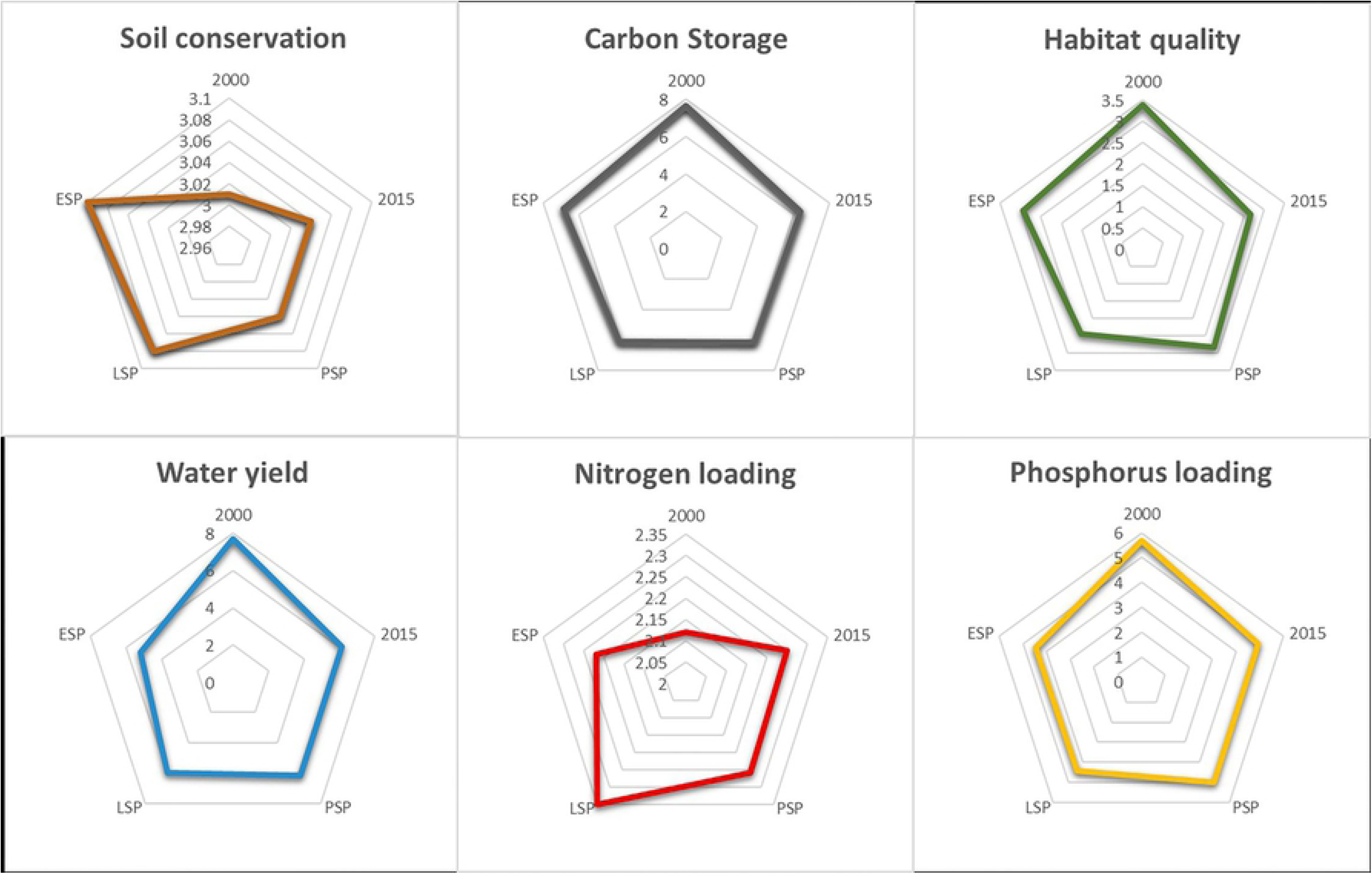
Actual and projected ESs supply from 2000 to 2030 in Nanjing.

We analyzed the trade-offs and synergies among various ESs to characterize the changes in ESs brought about by land use change between 2015 and 2030. We found that all ecosystem service pairs obtained statistically significant relationships except soil conservation and water production, phosphorus loading and phosphorus loading. There was a positive correlation between carbon storage and soil conservation, water production services and habitat quality, indicated that there was a synergistic effect among these ESs (Fig 5). Among them, the correlation between carbon storage and habitat quality was the strongest under EFP scenario, with a correlation coefficient of 0.957. At the same time, there was a positive correlation between carbon storage and nitrogen and phosphorus conservation, indicated that there was a trade-off between these ESs. We found that the trend of trade-off and synergy among various ESs was roughly the same throughout the study period, indicated that the change of land use types in Nanjing did not fundamentally change the trade-off relationship among various ESs.

**Fig. 5.**
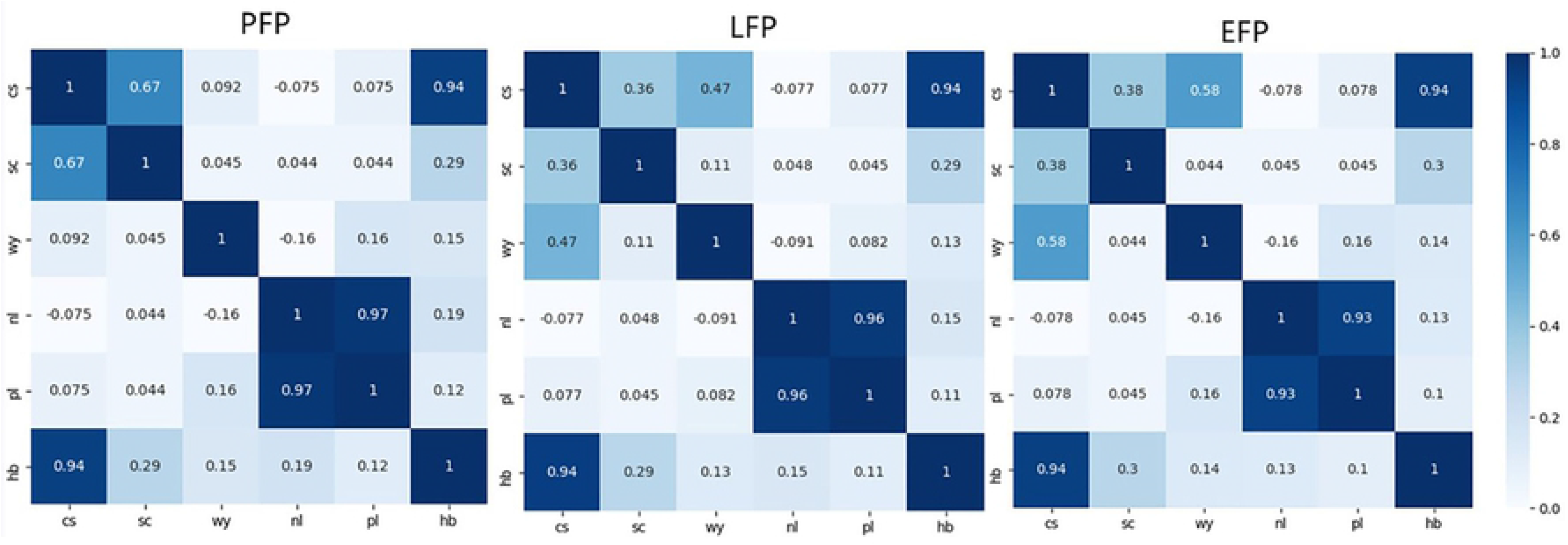
Correlation analysis between pairs of ESs in 2030 under various scenarios.

### Trade-offs and synergies among multiple ESs

In the PFP scenario, carbon storage, soil conservation capacity and habitat quality decreased, but water yield capacity and nitrogen and phosphorus loading increased, while carbon storage and habitat quality decreased when woodland was converted to urban land, water production capacity, soil retention capacity and nitrogen and phosphorus output increased. In the LFP scenario, carbon storage, water yield capacity, soil retention, habitat quality, and nitrogen and phosphorus loading all decreased when the cropland was converted to water area; when the cropland was converted to urban construction, carbon storage, habitat quality and nitrogen and phosphorus loading decreased, water yield capacity, soil retention capacity and nitrogen and phosphorus output capacity were increased. When unused land was converted to woodland, Carbon Storage, water production capacity, N/P loading decreased, while soil retention capacity and habitat quality improved. In the EFP scenario, carbon storage, soil conservation and habitat quality were increased while water yield capacity, nitrogen and phosphorus loading were decreased when cropland was converted to woodland; When cropland was converted to urban construction land; carbon storage, habitat quality and phosphorus loading were decreased; water yield capacity、 soil conservation capacity and nitrogen loading were increased; When the woodland was converted to urban construction land, carbon storage and habitat quality were decreased while water yield capacity, soil conservation and nitrogen and phosphorus loading were increased.

## Discussion

### Response of ESs to LUCC

Land-use pattern, land management policy, land-use planning and other factors are the main driving forces of ESs change. The complex relationship between ESs and LUCC has obvious regional and scale characteristics. Therefore, Therefore, it is essential to detect the relationship between them in different research areas, which can provide scientific basis for urban land use planning, ecosystem management and better human settlements construction. [44,45].

The result showed that the carbon storage, habitat quality and water yield capacity of Nanjing decreased significantly while the soil conservation capacity increased significantly from 2000 to 2015, the performance of the two indicators (N/P) of water purification was not the same. From different scenarios, the Carbon Storage and habitat quality scores in the EFP scenario increased [46].This because the large increase of the ecological land area such as woodland and grassland was in the Ecological space priority scenario, and it was from cropland, construction land and other land-use patterns with great man-made disturbance to woodland, grassland and other land-use patterns with small man-made disturbance. In the PFP scenario, the water yield was the highest because the farmland had a strong water yield capacity, and the model ignored factors such as snowfall and surface runoff. Water yield mainly comes from rainfall, and it could be through the vegetation evapotranspiration and surface infiltration, so there may be less vegetation cover, the more water yield; Soil conservation capacity was strongest in LFP scenario, indicated that both urban construction land and higher vegetation cover could enhance soil conservation capacity; the output of nitrogen in Nanjing increased from 2000 to 2015, but the output of phosphorus decreased, this indicated that the purification capacity of the city for nitrogen decreased and the purification capacity of the city for phosphorus increased during this period. In three different scenarios, phosphorus output was consistently declining, possibly because cropland was a source of growth for other land types. The output of nitrogen continued to increase in the PFP and EFP scenarios. Construction land has increased by 26.2 per cent and 15 per cent respectively since 2000, and this indicated construction land had a very high output of nitrogen. The output of nitrogen in LFP scenarios was reduced, even lower than the 2015 level, which indicated that the increase of ecological land can effectively achieve the goal of water purification. Meanwhile, we found that the conversion of unused land to other types of land can have beneficial effects, such as stabilizing sandy soil and reducing soil erosion.

Finally, we analyzed the trade-offs among various ESs and found that the trend of trade-offs among various ESs was approximately the same throughout the study period, it showed that the change of land-use types in cities can not fundamentally change the trade-offs among different types of ecosystem services, as found in other studies[47].Trade-off analysis has a potential role in land-use management environmental policy formulation and ecological civilization construction [48,49]; it enabled system makers to obtain a full picture of the potential impact of their policy on ESs trade-offs [50], and stroke a balance between intentional and unintentional ecological aftermaths [51].

### Strategies and implications

Sustainable Development is an important scientific goal to be fufilled by coping with the disordered exploitation of land resources, heavy ecological debt and deteriorating living environment [52]. City as a huge artificial integrated system, the operation of it inevitably brings the disturbance to the nature, this kind of disturbance leads to the trade-off and synergy of ESs. It is feasible to make clear the main function of the city and cooperate with a series of spatial development policies and control the disturbance of ESs within a certain range, and it is consistent with China’s spatial development strategy too. For example, for some ecological conservation areas or cities that provide key ESs, it is essential to minimize the disturbance of urban development to nature and implement the conversion of cropland to forests, grassland and wetlands, so as to further improve the quality of habitats. For areas with severe soil erosion and debris flow landslides [53], we need implement Afforestation to improve soil construction services and rainwater retention. For some cities with development potential, there may be a large number of ecological and productive function land convert to living function land, and lead to the decline of various ESs. How to achieve green development and balance the relationship between development and conservation is very important. For the areas with great development foundation and need to be optimized, the main function will no longer be the expansion of living function land, but the pursuit of higher habitat quality and less nitrogen and phosphorus loading.

For this case, we has implemented the dynamic simulation of land use change based on three functional scenarios. On the whole, the trend of build-up land under the three scenarios has been on the rise, which was in line with Nanjing’s basic vision of continuing to build new urbanization and modern cities. According to the requirements of “Nanjing City Master Plan (2018-2035)”, the total scale of build-up land at the end of the planning period need to be controlled at 2015 km^2^, a 14% increase in the scale compared with the previous period, which was similar to the growth area of build-up land in the PFP scenario. At the same time, among the three scenarios, the proportion of cropland in PFP scenario decreased the least, which was consistent with the policy of “Nanjing Land Use Master Plan 2006-2020”. Combined with other related research results, we put forward the following suggestions: (1) Focus on the impact of land use change on carbon storage services. Improving habitat quality and carbon storage by increasing urban green space is effective, but reducing CO_2_ emissions at source is a more effective way [54]. (2) Continue to stabilize agricultural production. Changes in land use resulting from rapid urbanization have reduced the overall quality and quantity of agricultural land [55].We should continue to promote strict and sustainable land use plans, such as the policy of “linking the increase of urban build-up land land use with the decrease of rural build-up land land use” [56]. (3) Further integrate the trade-offs of ESs with land-use decisions[57].There are significant differences in ESs among different LUCC types, and exist trade-off mechanism between ESs and environmental processes. (4) Determine the appropriate principal function policy. The relationship between LUCC and ESs is very complex, different decision-making scenarios have their own advantages and disadvantages. It is unscientific to select only one scenario. The real situation should be the superposition or combination of the scenarios based on the main function.

### Strengths and limitations

The use of spatial data and stakeholder engagement to assess multiple environments and societies was a fundamental feature of the Natural capital methodology philosophy [58]. The three scenarios implied not only the three main functions of urban development, but also the three different paradigms of urban development. In order to make the scenarios more realistic, we did not set the various scenarios as a “limiting” state. At the same time, we did not support the reckless classification of the six land types into three types of land use: production, living, ecology, although some research results did before. What we advocated was the classification based on the main function, which means give the main function of urban development, but did not exclude the existence of other functions. For example, the scenario in which ecological protection was the main function did not mean that the encroachment of production space and living space on ecological space was absolutely prohibited. Based on the comprehensive consideration of various situations, policies and natural background, the main function-oriented land classification was more in line with the transformation trend of China’s land spatial planning.

At the same time, the InVEST model can be used to better evaluate the spatial and temporal distribution of ecosystems and their changes in land use/cover scenarios, and to provide spatial planning recommendations for land use decisions [59,60]. However, due to the lack of monitoring data, model defects and evaluation indicators, there were some uncertainties in the results of the whole study area. For example, the Nutrient export model of InVEST had limited input parameters and did not take into account the chemical interactions between nutrients, so there must be a discrepancy between the results of the model and the actual situation. The soil conservation model of InVEST derive soil conservation amount from the difference between potential and actual soil erosion. The model only considers the erosion in flow, and did not consider the sediment background, such as the factors of Rivulet, bank, dissolution of flowing water, etc. therefore, there were some differences between the results of the model and the study that took these factors into account [61]. Nevertheless, the results of the Invest Model as a reaction to an overall trend were acceptable [62]. The MODEL provided the information of ESs and still reflected the contrast of time series and the characteristics of space-time distribution. It had scientific guiding significance for locating the priority areas of ESs, reasonable ecological protection and planning practice [63].

## Conclusions

In this study, we assessed the response of LUCC and ESs in Nanjing from 2000 to 2015, and found that carbon storage, habitat quality and water yield capacity in Nanjing declined by 17%, 21% and 20%, respectively, soil and water conservation capacity remains stable. We studied three functional scenarios to simulate the response of LUCC and ESs to different functional scenarios from 2015 to 2030. According to our assessment, from 2000 to 2015, carbon storage, habitat quality and water yield capacity in Nanjing decreased significantly, soil conservation increased slightly, and the performance of the two indicators (N/P) for water purification was not consistent. From different scenarios, carbon storage and habitat quality were the highest in EFP, water yield was the highest in PFP and soil conservation was the highest in LFP. It also found that from 2000 to 2015, Nanjing’s purification capacity for nitrogen decreased, but its purification capacity for phosphorus increased. In three different scenarios, phosphorus output was in a decreasing trend, while nitrogen output was in a decreasing trend in LFP scenarios. We analyzed the trade-offs and synergies of ESs in various functional scenarios, and notice that the trade-offs among different types of ESs tended to be the same throughout the whole period, it has showed that the change of land use types in cities did not fundamentally change the trend of trade-off and synergy among various ESs. We found that different functional scenarios have different impacts on ESs, and we believed that the determination of the main function of the city was the first condition to judge the applicability of scenario.

## Acknowledgements

Thanks to the data support of National Earth system science data center and Resources and Environment Data Cloud Platform of Chinese Academy of Sciences.

## Supporting information

**S1 Fig.**
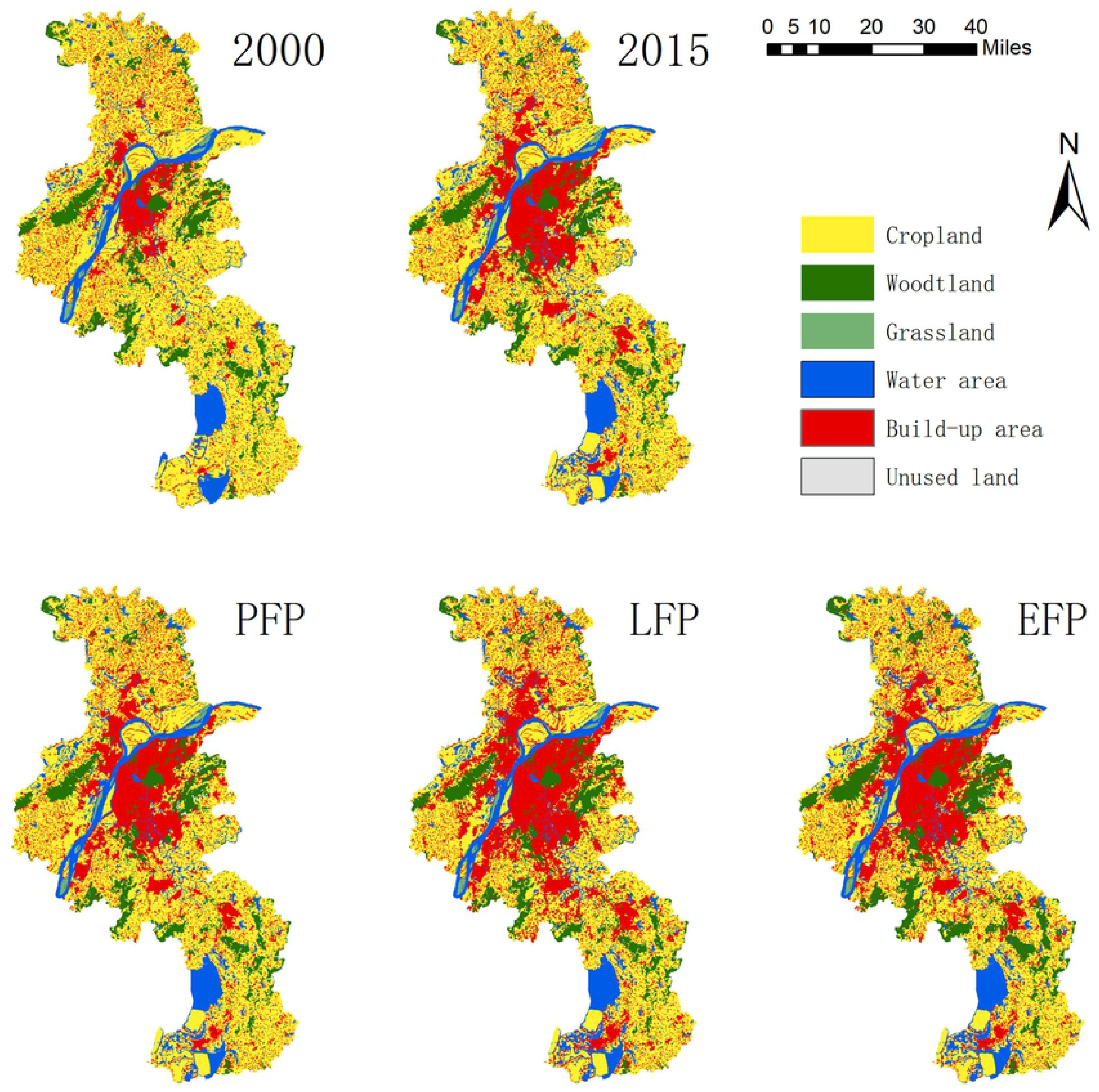
Land use change in Nanjing from 2000 to 2030.

**S1 Table Cost matrix of land use conversion under different scenarios in Nanjing**

**S2 Table The determination of elastic parameters of land use types under different scenarios**

**S3 Table Biomass carbon density of different land use types in Nanjing (Mg/ha)**

**S4 Table Biophysical coefficient table of water yield module**

**S5 Table Table of biophysical coefficients of soil conservation module**

**S6 Table Biophysical table of nutrient delivery ratio**

**S7 Table Habitat types and their sensitivity to threats**

